# Scalable design of repeat protein structural dynamics via probabilistic coarse-grained models

**DOI:** 10.1101/2024.03.13.584748

**Authors:** Seeralan Sarvaharman, Timon E. Neary, Thomas E. Gorochowski, Fabio Parmeggiani

**Author notes:** Correspondence should be addressed to S.S., T.E.G. and F.P. These authors should be considered as joint senior author with equal contribution (Thomas E. Gorochowski, Fabio Parmeggiani).

## Abstract

Computational protein design has emerged as a powerful tool for creating proteins with novel functionalities. However, most existing methods ignore structural dynamics even though they are known to play a central role in many protein functions. Furthermore, methods like molecular dynamics that are able to simulate protein movements are computationally demanding and do not scale for the design of even moderately sized proteins. Here, we develop a probabilistic coarse-grained model to overcome these limitations and support the design of the structural dynamics of modular repeat proteins. Our model allows us to rapidly calculate the probability distribution of structural conformations of large modular proteins, enabling efficient screening of design candidates based on features of their dynamics. We demonstrate this capability by exploring the design landscape of 4–6 module repeat proteins. We assess the flexibility, curvature and multi-state potential of over 65,000 protein variants and identify the roles that particular modules play in controlling these features. Although our focus here is on protein design, the methods developed are easily generalised to any modular structure (e.g., DNA origami), offering a means to incorporate dynamics into diverse biological design workflows.

## INTRODUCTION

The structural dynamics of proteins play a crucial role in their function and contribution to a wide variety of biomolecular processes ^1^. Examples range from the active transport of molecules ^2^, to the sensing of stimuli ^3^. In the field of synthetic biology, computational protein design has emerged as a powerful tool for creating new proteins with desired functionalities. It has been used to support the design of florescence activating proteins ^4^, triose-phosphate isomerase (TIM) barrels ^5^, proteins that can triggering immune responses ^6^, enzymes for catalysis ^7^ and protein switches ^8^, to name but a few. However, while the central role of protein structural dynamics is well known ^9^, when it comes to engineering *de novo* proteins, their dynamics for the most part have been neglected.

The most detailed predictions of protein dynamics are generated using molecular dynamics (MD) simulations ^10–12^. These provide atomistic detail and can capture the complex motions that proteins exhibit. Unfortunately, MD simulations are still time-consuming to perform and require substantial computational resources to be performed at scale, especially when simulating large proteins. Moreover, to achieve accurate prediction of dynamics, extensive sampling and long simulation times are necessary, limiting their application for selecting candidate designs from large variant libraries or to support rapid iterative design cycles. Some of these difficulties have been partially addressed by exploiting hybrid modelling approaches ^13,14^, however, scalability issues remain.

To overcome the computational limitations of molecular simulations, coarse-grained models have been developed ^15–17^. These use simplified representations of proteins, typically abstracting multiple atoms as a single interaction site. This reduces the degrees of freedom in the model and enables simulations over much longer timescales. Coarse-grained models in some cases have been found to capture the essential structural and dynamical features needed for design tasks, while significantly reducing the computational demands ^18,19^.

A common coarse-grained approach for the prediction of protein dynamics is the elastic network model (ENM) ^20,21^. ENMs approximate a protein structure as a network of interconnected springs, where each spring represents an interaction between two residues ^22^. The model captures the collective motions of the protein by considering the harmonic vibrations around the equilibrium positions ^23^. ENMs provide a simplified representation of protein dynamics and have been successful in modelling global, low-frequency motions and functionally important motions away from thermal fluctuations. However, they do not always fully capture higher frequency movements or the details of local interactions that result from non-collective motions ^24^. Furthermore, the number of spring constants that must initially be fit changes depending on the number of amino acids present. This makes it difficult to rapidly evaluate similar candidates that differ in length, or whose model parameters are drastically different. Therefore, ENM models are typically unsuitable for iterative design workflows, where many diverse designs needs to be quickly evaluated at each cycle.

In this work, we aim to overcome these limitations and strike a balance between the computational efficiency of a coarse-grained representation and the ability for it to capture key protein dynamics. We focus on the simulation of tandem repeat protein domains, as they can reach hundreds of amino acids in size, resulting in large global motions as well as confined local motions ^25^. Repeat proteins are found widely in nature ^26,27^ and perform diverse biological functions ^28–30^. They are characterised by the presence of repetitive structural motifs/modules, which offers several advantages for modelling and *de novo* protein design. First, their modular nature greatly simplifies the prediction of tertiary structures, enabling the more rational design of novel sequences based on known repeat modules ^31^ and the combination of different repeats ^32^. Second, and most importantly, repeat proteins exhibit more predictable structural dynamics within each module ^33–35^, making them a useful platform upon which dynamics can be effectively modelled and exploited in the design process ^36^. By capitalising on the repetitive architecture of these proteins, we are able to construct a coarse-grained model that can efficiently propagate expected structural dynamics through the chain of modules making up a protein (**Figure 1**). We demonstrate how the speed of our model allows us to predict, score and extract profiles of protein dynamics in a modular design landscape, offering a means to quickly and reliably discover candidates with desired structural and dynamical characteristics. In addition, we show how this comprehensive view of a design space can provide valuable information regarding the flexibility and responsiveness of candidate modules to the sequence context, and offer insights into how specific modules are likely to affect overall features of a larger repeat protein. As protein design moves towards applications that require the careful crafting of conformational changes in protein structure, our model provides a means to assess such features and support engineering workflows that place dynamics at the forefront.

**Figure 1:**
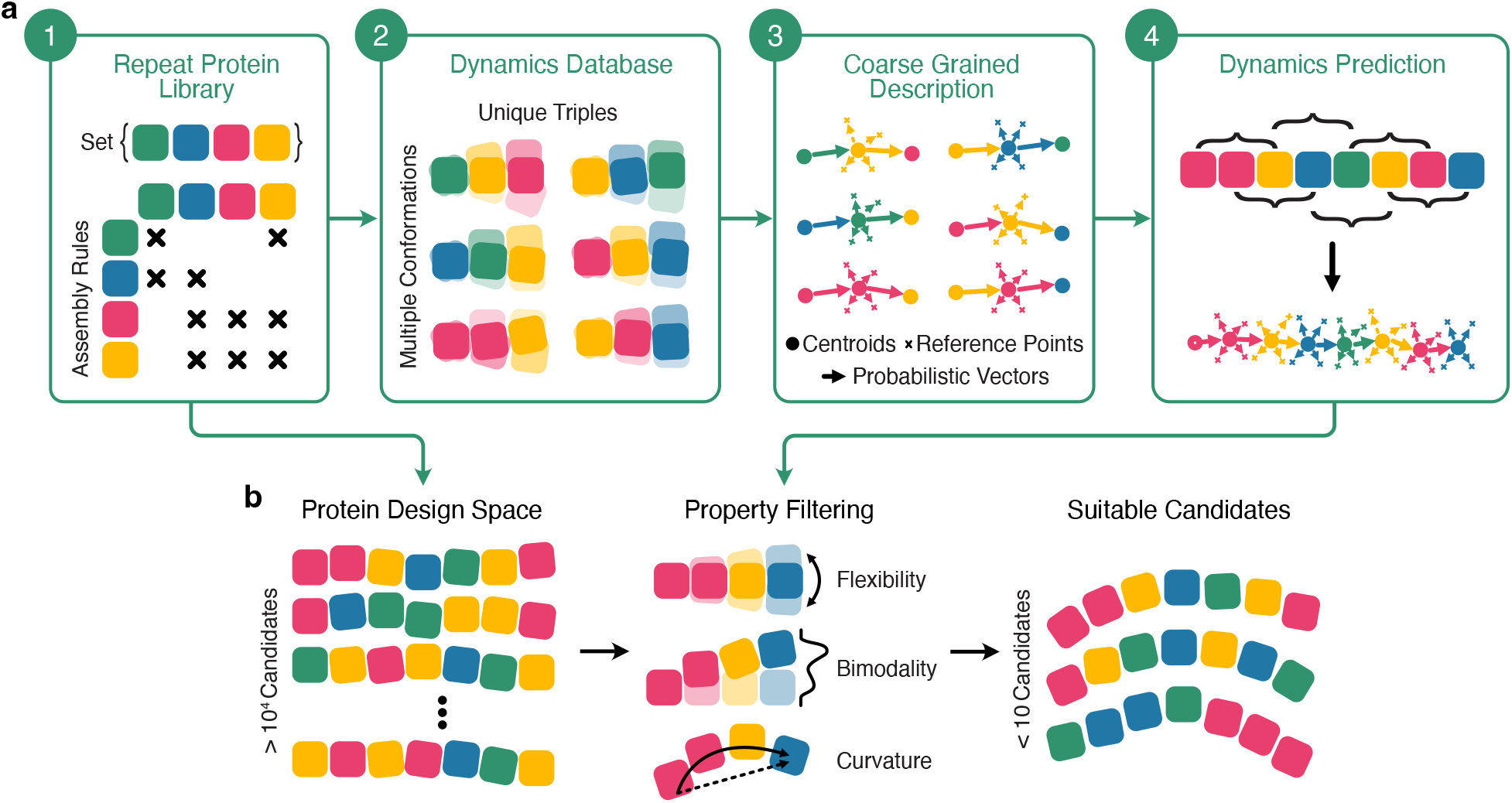
Overview of the coarse-grained model for the structural and dynamical design of modular proteins. (**a**) Workflow for generating a coarse-grained model of protein structural dynamics. The model relies on an underlying repeat protein library that contains well defined protein modules and rules for how these can be assembled (Step 1). Using this information, a dynamics database of all three module proteins capturing the relative movements between all atoms in the protein is generated (Step 2). This can be via detailed simulations (e.g., using Rosetta) or derived from experimental data. The resultant dynamics database is then used to build coarse-grained descriptions of each triplet of modules, defining centroids for each module and an arbitrary number of reference points, e.g., the end points of alpha helices (Step 3). Finally, these coarse-grained descriptions are stitched together to enable the efficient propagation of movements of larger proteins build from the repeat protein library (Step 4). (**b**) The coarse-grained model can be used to accelerate the design of proteins built using the repeat protein library. A typical library defines a vast potential design space that would be impossible to exhaustively search. The speed of the coarse-grained model allows for large regions of this space to probed (e.g., millions of designs) and key structural and dynamical properties measured (e.g., flexibility or potential for multi-state dynamics). This information can be used to guide future areas to explore and the properties of the simulated designs can be filtered on properties that are essential to the desired function of the protein. Using this approach a targeted set of designs that can be feasibly built is output for detailed experimental testing.

## RESULTS

### Coarse-grained model of repeat protein dynamics

The input to our model is a repeat protein library ^37^ consisting of a set of protein modules (with each module comprising at least two repeats) and a connectivity matrix that defines the rules for assembling larger constructs, i.e., for a given module, which other modules are compatible and can be directly connected (**Figure 1a**, step 1). Note that these rules do not necessarily commute, such that “module B can follow module A” does not imply “module A can follow module B”. This library defines the overall accessible design space. However, it can be extended at any time by adding further modules and connectivity rules. We chose to use an existing repeat protein library that contains 34 modules that are on average 180 Å long, covering a wide range of structures ^38,39^.

In addition to the repeat protein library, we also require a dynamics database that can be used by the model to predict the movement of larger multi-module proteins (**Figure 1a**, step 2). The database consists of a large number of conformational snapshots for all possible combinations of three-module constructs. For our library, this equates to a total of 644 unique constructs that were 320 to 794 amino acids long, and 100 conformational snapshots for each. These snapshots can be identified with minima in the rough energy landscape of the protein, and the movement of the protein is driven by thermal fluctuations, which on a slow enough timescale cause jumps from one minima to another. These snapshots therefore allow us to infer the steady-state dynamics of proteins using a probability distribution of atomistic positions.

Obtaining this information via experimental methods such as hydrogen-deuterium exchange via Nuclear Magnetic Resonance (NMR) or mass spectrometry (HDX-MS), is infeasible due to the total number of unique constructs in our library and the size of the molecules. Using computational methods like molecular dynamics (MD), simulations were also not feasible due to the size of the proteins. We therefore chose to use conformational snapshots generated using the relax protocol from the Rosetta modelling suite ^40^ (**Methods**). We based our database on three module constructs due to this being the smallest number of modules where contextual effect of different neighbours on a given module can be observed. While it is possible to utilise higher-order contextual effects by using constructs with more than three modules, obtaining snapshots becomes computationally challenging due to the larger number of constructs that must be assessed and the increased number of residues per construct.

For the model to efficiently use the information within the dynamics database, we generated coarse-grained descriptions that simplify the propagation of dynamical information throughout larger multi-module proteins (**Figure 1a**, step 3). We chose key anchor points, specifically the ends of *α*-helices, to capture the orientation of the module and then define the mean of these key anchor points as the centroid of the module. The locations of the centroids and the anchor points change depending on the conformation. Therefore, we use the conformational landscape in the dynamics database to build a probability density estimate for the locations of the centroids and their anchor points. As these descriptions are analogous to position vectors when the dynamics are neglected, we refer to them as probabilistic vectors. For each unique triplet of modules (*a · b · c*), the coarse-grained model is then generated as follows. We first perform a rigid body transformation on all of the conformations such that the centroid of the central module *b* (***c***_*b*_) is located at the origin (0), and rotated appropriately such that any rotational symmetry from the placement of modules is removed. Using the conformational data, we then calculate the probability distribution *P*(***c***_*a*_|***c***_*b*_ = **0**) and *P*(***c***_*c*_|***c***_*b*_ = **0**), which captures the steady-state occupation probability density of module *a* relative to module *b*, and *c* relative to module *b*, respectively. For module *b*, we can also track an arbitrary number of reference points ***r***_*i*_ (e.g., start and ends of alpha helices) via *P*(***r***_*i*_|***c***_*b*_ = **0**), which describes the steady-state occupation probability density of the *i*^th^ reference point in module *b* relative to the centroid of module *b*. These probability density functions are estimated by fitting a Gaussian mixture that can be stored efficiently using the parameters of the mixture (i.e., means, covariances and weights of the constituent components). We precompute these parameters for all of the module triplets and use them for efficiently estimating the dynamics of larger constructs.

Finally, to predict the dynamics of arbitrarily sized modular proteins, we use this description to estimate the steady-state occupation probability density of the *k*^th^ centroid in a many-module construct by identifying the constituting triplets that make up the larger construct, and perform the convolutions

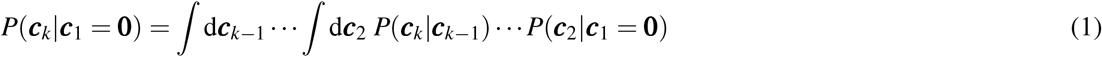

when *k ≥* 3 and where we condition the location of the centroid of the first module to be at the origin (**Figure 1a**, step 4). To illustrate the procedure, we can consider a three module construct, labelled 1, 2 and 3. With the centroid of the first module ***c***_1_ centred at the origin, *P*(***c***_2_|***c***_1_ = **0**) describes the position of the second centroid ***c***_2_ relative to the first. Similarly, *P*(***c***_3_|***c***_2_ = **0**) describes the position of the third centroid ***c***_3_ relative to a fixed ***c***_2_ at the origin. In order to obtain the distribution of the position of ***c***_3_ relative to the the fixed ***c***_1_ at the origin, one needs to perform a convolution of the distributions. The probability density estimate for the reference points in the *k*^th^ module can also then be computed via:

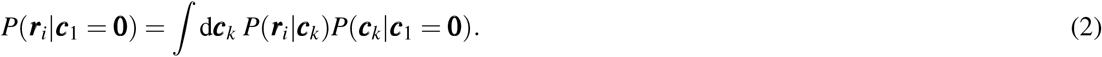

Together, these equations allow us to propagate the local movements captured in our dynamics database through to larger constructs and estimate diverse structural dynamics from the centroids and reference points.

This algorithm was implemented in a package called Dynamo (**Data Availability**). Dynamo is a native Python library written in Rust to ensure reliable and high-performance model generation and simulation. It also includes additional helper functions to simplify the creation of the dynamics database, the ability to define complex multi-module constructs beyond chains (e.g., star-like proteins), visualisation tools to better understand the inferred protein dynamics, and the ability to export data in standard Protein Data Bank (PDB) format for use in other tools.

### Model validation

To verify the accuracy of our model, we assessed the differences between the dynamics predicted from our coarse-grained model with those extracted from the conformations obtained from the Rosetta relax protocol. We chose to consider a diverse set of 14 modular proteins where 10 were homogeneous containing 9 modules of the same type, while 4 were heterogeneous containing 4 modules of different types. These proteins ranged in size from 701 to 840 amino acids long.

To compare the qualitative agreement between our model and the Rosetta relax data, we computed the displacements, ***r***, between the specific centroids of the conformational samples and the mean centroid location across them all, and compared the distribution of its magnitude |***r***|. This distribution captures both the overall magnitude of any movement, as well as its shape. For example, if the centroid was distributed on the surface of a sphere of radius *R*, then the distribution of |***r***|, would reduce to the Dirac-delta function *P*(|***r***|) = *δ* (|***r***| *− R*). As the point cloud of ***r*** becomes more complex in shape, the distribution *P*(|***r***|) will exhibit more complex features.

For each modular protein construct, we plotted the distribution *P*(|***r***|), for the 3^rd^, 6^th^, and 9^th^ centroids for the homogeneous constructs, while the 2^nd^, 3^rd^, and 4^th^ centroids for the heterogeneous constructs (**Figure 2a**). In most cases, we found model predictions agreed well with the Rosetta relax data (i.e., the modes of the distribution coincided with each other). The main exception was the D4 construct, where the model predicted more movement than expected. A potential reason for this disagreement could be the larger number of rare conformations of the D4-D4-D4 triplet in the dynamics database used to parameterise the model. This would result in a wider exploration of the conformational landscapes and cause the model to infer larger movements that get propagated through the entire protein. For the H4 construct, we also overestimated the dynamics. However, the differences are smaller than those of the D4 construct.

**Figure 2:**
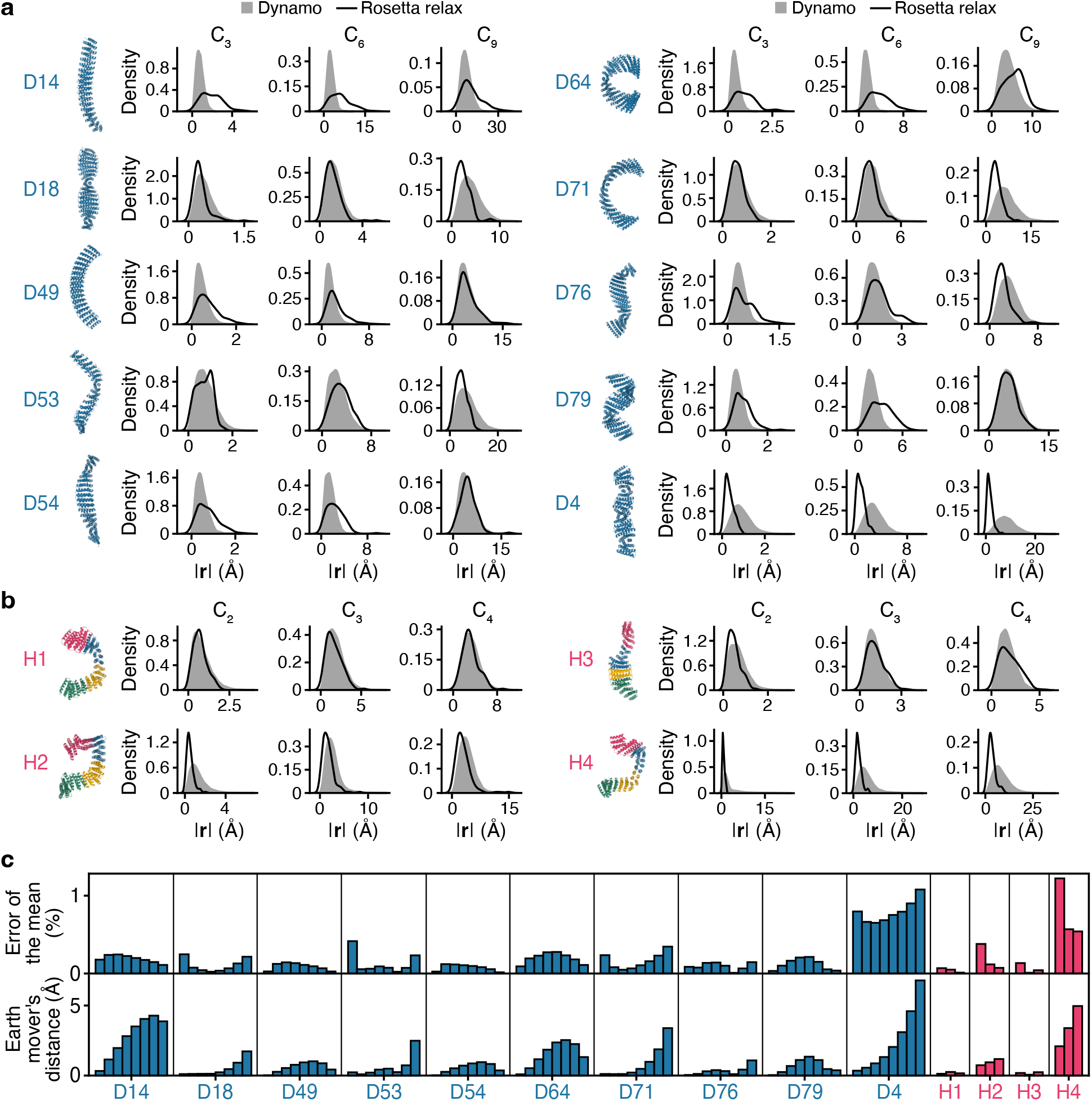
Comparison of conformational data obtained using the Rosetta relax protocol and our model (Dynamo). (**a**) Distributions capturing the fluctuations in the position of centroids (C_*x*_) relative to their respective mean. Each centroid has a mean position, and the displacement from this mean and samples of centroid positions obtained from our model (grey filled distribution) or through Rosetta relax (solid black line) is given by **r**. Distributions shown for ten homogeneous 9-module constructs. A molecular visualisation of the construct is shown on the right of each plot. (**b**) Similar distributions as described in panel (a) for four heterogeneous 4-module constructs. The modules in the heterogeneous constructs are: H1 = D14 j1 D14x4 188; H2 = D14 j1 D18x4 31; H3 = D49 j1 D49x4 49; H4 = D79 j1 D14x4 131. (**c**) Comparison between the probability densities in panels (a) and (b) between the model and the Rosetta relax data for each centroid: (top) percentage error of the mean of the distributions, (bottom) a full distribution comparison using the earth mover’s distance. Bars correspond to centroids C_2_ to C_9_ (left to right) for the 9-module homogeneous constructs and C_2_ to C_4_ (left to right) for the 4-module heterogeneous constructs.

To compare the distributions further, we computed the percentage error in the means of the distributions for each of the centroids (**Figure 2b**, top panel). Again, we found that the largest differences between the model and Rosetta relax data was for the D4 construct. However, the average error across all constructs and centroids was only 0.24% which further provides evidence for the accuracy of the inferred movements of the proteins. To better assess the similarity in the shape of the distributions, we also calculated the Earth mover’s distance for each centroid in every protein construct (**Figure 2b**, bottom panel). We found that the average Earth mover’s distance across all centroids and constructs was only 0.95 Å, suggesting good quantitative agreement despite the coarseness of our the model.

### Visualisation of protein structural dynamics

During model validation, it became clear that it was difficult to assess subtle differences in protein dynamics due to the need to observe both structural and movement features of the data simultaneously. To help overcome this, we developed a new visualisation technique that captures the major dynamical features of each module in the context of the core alpha-helices making up the protein (**Figure 3**). The visualisation is created by first extracting the steady-state distributions of each centroid within a construct and calculating the covariances in their movement. For each module covariance, the direction and magnitude of the principal variances can be described as three mutually orthogonal vectors. These can then be used to generate ‘fins’, spanning either side of the mean centroid position and parallel to the principal variances. To aid comparison, we colour the fin at a particular point in relation to the total variance in the distribution of centroid locations, which corresponds to the magnitude of any movement. We also overlay each helix as a semi-transparent grey cylinder to further portray the protein’s underlying structure. Using this visualisation technique, we could place the errors in the context of the protein size and shape and clearly see that our coarse-grained model was able to accurately capture the specific magnitude and direction of protein dynamics when compared to the Rosetta relax data (**Figure 3**).

**Figure 3:**
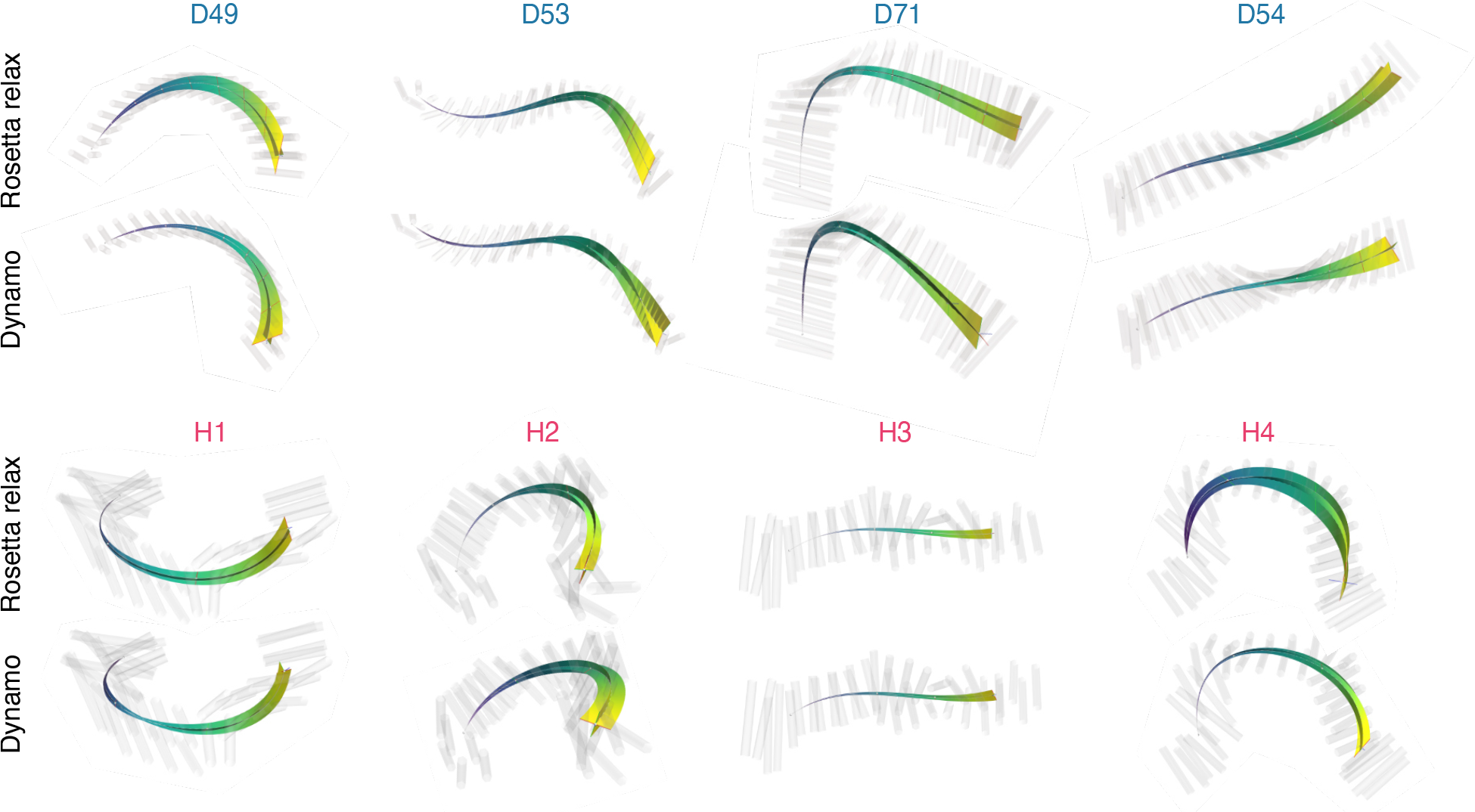
Visualisation of the structural dynamics of several modular protein designs. Data shown for the Rosetta relax protocol (top) compared to our model (bottom). The two fins are parallel to the two directions with the most movement and envelope the 95^th^ percentile of the centroid density distributions, while the colour and width of the fins correspond to the largest movement (dark blue to yellow denoting small to large movements, respectively). The alpha helices are represented by grey semi-transparent cylinders. The modules in the heterogeneous constructs are: H1 = D14 j1 D14x4 188; H2 = D14 j1 D18x4 31; H3 = D49 j1 D49x4 49; H4 = D79 j1 D14x4 131.

### Mapping out the design landscape of a repeat protein library

The efficiency of our model enables us to predict the structural dynamics of large modular protein design spaces. To demonstrate this, we considered all possible 4-module protein designs using our modular protein database. We generated the structural dynamics of each of the 2,978 designs within this space and quantified two key features that covered the structural geometry and dynamics of each design. The first was the protein curvature *α*, defined as the ratio between first and last centroid distance and corresponding arc length. The second was the flexibility *β*, which captures the overall extent of the dynamics (see details in methods). Plotting the density of designs in relation to these features provided us with a uniquely complete picture of the available characteristics of all 4-module proteins (**Figure 4a**).

**Figure 4:**
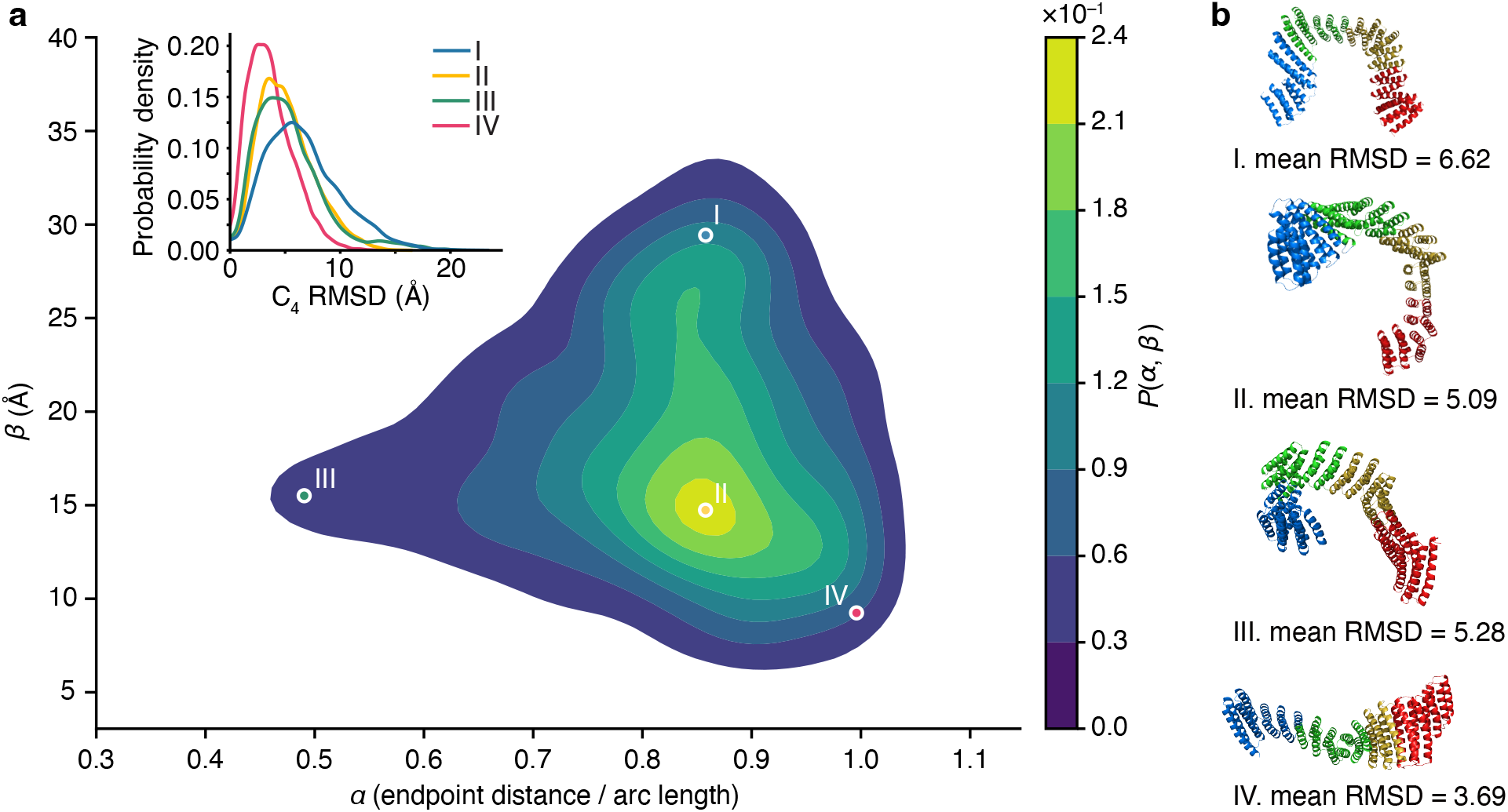
Exploring the design landscape of all 4-module protein chains containing 2,978 unique designs. (**a**) Smoothed density plot of all 4-module protein chains from our model. The position of four selected designs is indicated. Note that 0 *≤ α ≤* 1, but kernel density estimates can give non zero probability outside of this range, for clarity of visualisation we have chosen not remove the small region of probability estimate for *α >* 1. Insert shows the probability distribution of the root mean squared deviation (RMSD) of the final module in the chain (C_4_) for the four highlighted designs (I–IV), as calculated from full length Rosetta relax runs. This distribution captures the general range of dynamic movements the module experiences and correlates with the flexibility (*β*). (**b**) Structural visualisations of specific protein designs highlighted in panel a. Individual modules denoted in different colours from N-(blue) to C-terminus (red).

Within the design space, we selected four examples with different properties and verified their qualitative agreement with data obtained from the Rosetta relax protocol. Our model predicted design I to be the most flexible (highest *β*) and design IV the least (lowest *β*). This was verified by the mean Root Mean Squared Deviation (RMSD) of the final centroid being the highest and lowest, respectively, and by distributions with the largest and narrowest distributions (**Figure 4a**, inset). Moreover, the model also captured the fact that designs II and III have very similar flexibility profiles corresponding to similar values of *β* and *C*_4_ RMSD distributions (**Figure 4a**, inset), whilst having very different sinuosity (**Figure 4b**).

### Proteins with multi-state potential

The ability for a protein to adopt several distinct conformational states plays a crucial role in a wide variety of cellular processes spanning, the function of molecular motors ^41^ to improved catalysis ^42^. From a design perspective, being able to suggest candidate protein designs with an inherent propensity for multi-state conformations would be valuable as a starting point for switchable and dynamic protein functions. While both globular and non-globular proteins can exhibit multi-state dynamics, repeat proteins represent a highly predictable and versatile platform to design multi-state systems. Due to the localised interactions, multi-state dynamics exhibited by a repeat protein can emerge from the behaviour of the individual modules.

We capitalised on this feature and developed a method to score candidate protein designs based on their potential for multi-state dynamics. Because multi-state dynamics are manifested as multi-modal probability distributions in the centroid positions, we could quantify the multi-modality of a construct by considering the probability density of the centroid of the final module. To do this, we first sampled from its probability density estimate. These samples were then split into training and testing sets using a *k*-fold validation scheme (*k* = 5). We used the training set to fit a Gaussian mixture model with the number of components, *n*, ranging from 1 to 5. For each fit, we obtained a score given by the likelihood of the appropriate test sets, which was then normalised appropriately to construct a probability mass function over the number of components, *p*(*n*). The mode of this distribution corresponded to the modes of the centroid density, *γ*_1_. To score this aspect (i.e. presence of distinct multiple-modes), we then computed the reciprocal of the entropy of the probability mass function:

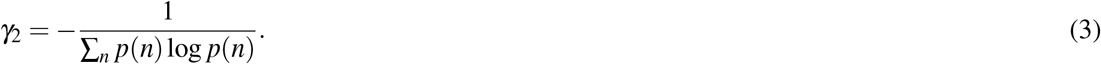

Low values of *γ*_2_ indicate a low confidence in the number of modes of the probability distribution of centroid, whereas high values a high confidence. Using this scoring we could rank candidates both on the number of modes present in their distribution and our confidence in this property.

To demonstrate this approach, we again considered all 4-module constructs and screened them for potential multi-state dynamics. Of our eight top scoring candidates, several showed bi-stable or multi-stable dynamics with H10 and H11 being the most promising (**Figure 5**). Many displayed very wide distributions with no clearly separated modes. While these candidates may not exhibit strong multi-state behaviour, the large range of movement (60 Å for H5) would provide an ideal starting point for establishing multi-stable dynamics using external stimuli, such as an additional peptide chain ^43^.

**Figure 5:**
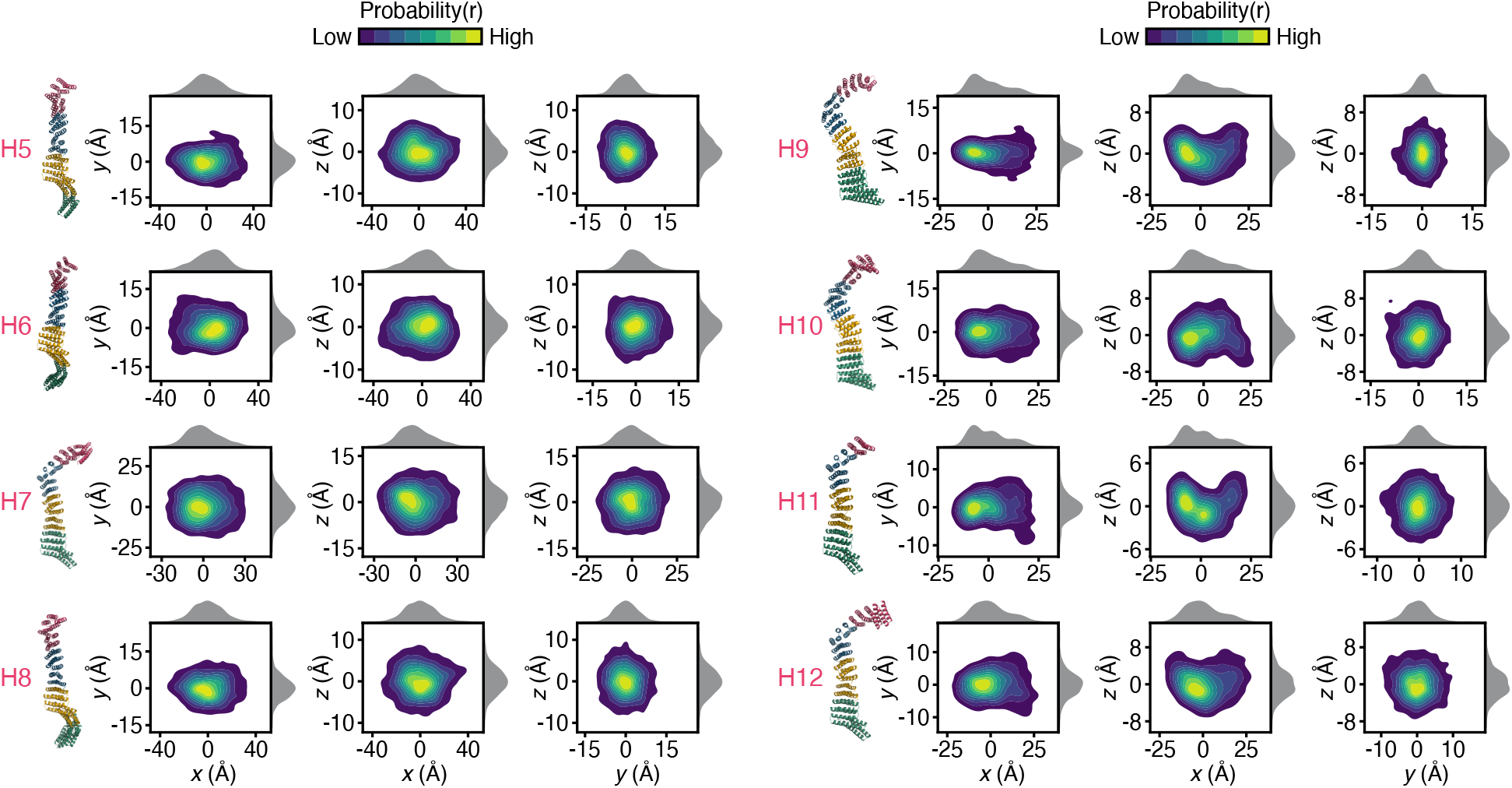
Exploring potential multi-stable behaviour of 4-module constructs. The eight most highly-ranked constructs after scoring each on their potential for multi-stable behaviour. The position of the centroid of the last module, *r*, is visualised as a probability distribution projected onto orthogonal planes. Each distribution is built by sampling 10^6^ centroid positions from the model. The modules in the heterogenous constructs are: H5 = D14_j1_D14x4_239; H6 = D14_j1_D14x4_241; H7 = D79_j2_D14x4_117; H8 = D14_j1_D14x4_240; H9 = D18_j1_D14x4_117; H10 = D14_j1_D14x4_117; H11 = D14x4_117; H12 = D49_j1_D14x4_117.

### The effect of specific modules on construct rigidity

The ability to generate the structural dynamics for entire modular protein design spaces, offers the ability to unravel the potential roles that individual modules might play more broadly across many different designs. A key feature that often needs to be controlled when designing *de novo* proteins is the rigidity of the final design. This can be quantified by calculating *β*, which captures the likely overall movement of a protein from the probabilistic conformational data (**Methods**). Given that the overall rigidity of a construct is determined by the modules present (e.g., some modules might stabilise the module while others might introduce more flexibility), it is possible to uncover the influence each module has when they are part of a larger construct. To do this, for a given module in an *n* module construct, all possible constructs can be separated into *n* + 1 subsets based on the frequency of that module within the construct (**Figure 6a**, left panel). For each of these subsets, a probability density function can be built based on the constructs present. To compare how the properties of these distributions change with the different counts of a particular module, we can then visualise the means and variances of the distributions in a two-dimensional parameter space plot (**Figure 6a**, right panel). The mean of the distribution (Mean *β*) denotes the average flexibility of the population, while the variance of the distribution (Var *β*) quantifies the heterogeneity. As the occurrences of a specific module increases, the trajectory in this parameter space determines the role that the module plays within the larger constructs. There are four key types of behaviour: (i) universally stabilising, (ii) universally destabilising, (iii) contextually stabilising, and (iv) contextually destabilising. Increasing the counts of a stabilising module in a construct reduces its flexibility which causes the mean of the distribution *P*^*n*^(*β*) to decrease. When this stabilisation effect is universal, adding more of the given module in a construct drives down the dynamics regardless of the local context where the module is being added, or the presence of other modules which in turn causes the variance of *β* to also decrease. Conversely, when the stabilising is contextual, the local context and combination of other modules present play more of an important role in the dynamics of the construct. In other words, the flexibility sub-population remains highly heterogeneous with increases in module counts which yields less reduction in the variance of *β* when compared with the analogue of the whole population. On the other hand, increasing the counts of a destabilising module causes the flexibility of the constructs to also increase. When the effect is independent of other contextual factors then it is called strongly destabilising yielding decreases in the variance of *β*. Whereas the weakly destabilising are those that increase the flexibility of the construct, but only within specific contexts. It should be noted that while we have here focused on flexibility/rigidity of the constructs, this types of approach can be used for any feature that can be calculated from the structural dynamics.

**Figure 6:**
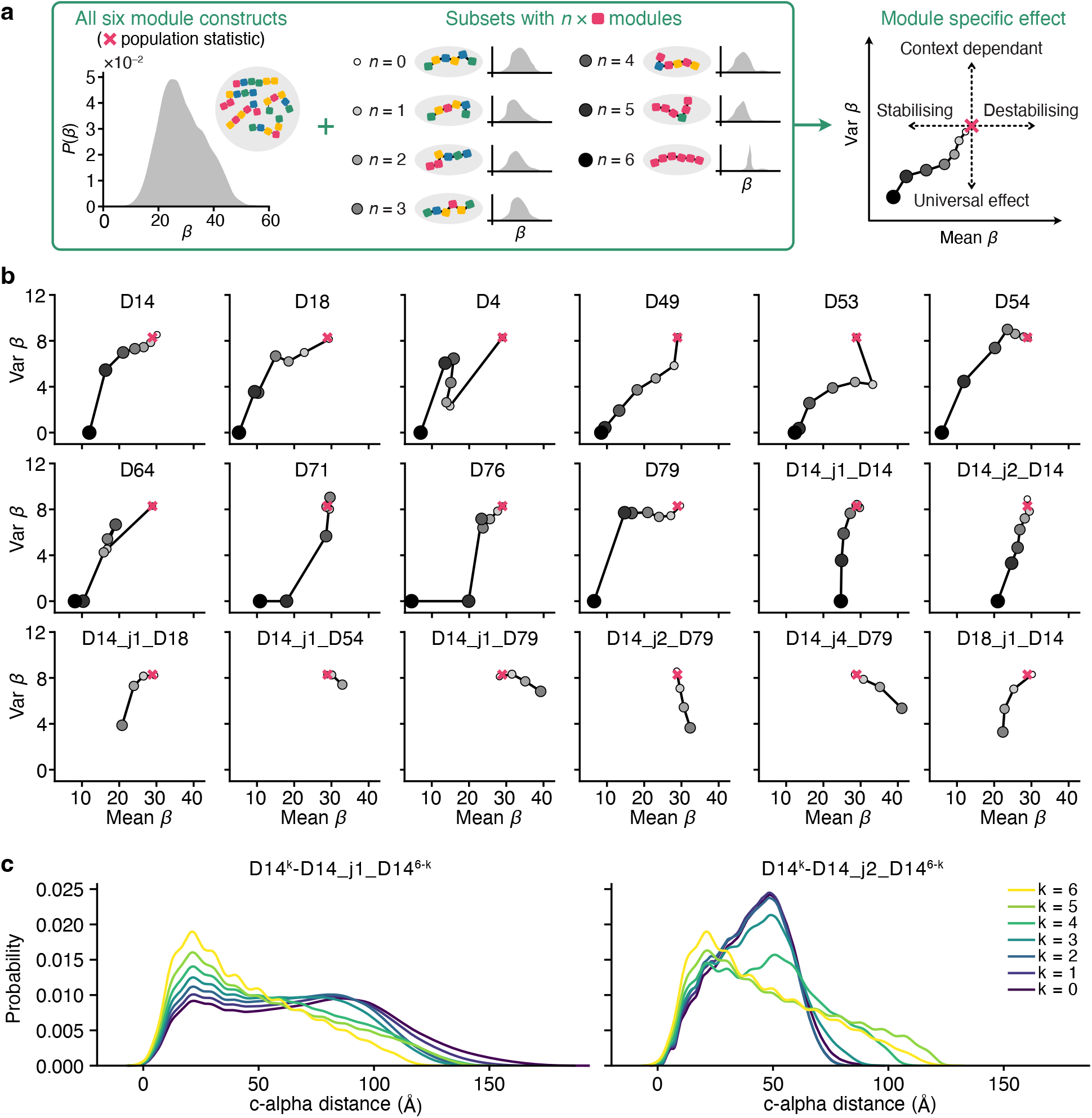
Analysing the role of each module on the flexibility of 6-module constructs. (**a**) The module specific effect is summarised by a plot showing the mean and variance of *β* probability distributions, with points for constructs containing 0 to 6 instances of the modules of interest (small white filled to large black filled circle). A summary statistic is plotted (red cross) of the entire set of all 6-module designs. Changes in the mean and variance of *β* as the number of instances of a module increase relate to stabilising or destabilising effects that are either context dependant or universal for all other modules. (**b**) Module specific effect plots for each module in every 6-module construct. (**c**) Probability density of all-to-all pairwise distances of alpha carbons in a construct containing a combination of D14 and D14_j1_D14 or D14_j2_D14, normalised to the maximum distance of a pure D14 construct (177 Å). *k* indicates the number of D14 modules in the protein.

To quantify the role that modules had in relation to protein rigidity, we generated the structural dynamics for all possible 6-module constructs and assessed the parameter space plots for 18 modules in our database (**Figure 6b**). These showed a broad range of behaviours across the modules. Most prominent was a universal stabilising effect, displayed by many of the modules and most prominently by D49, D54, D18 and D14. D79 showed a less universal stabilising effect, while several modules displayed non-uniform behaviours (e.g., D4, D53 and D64). We also found that 4 of the modules were universally destabilising (D14 j1 D54, D14 j1 D79, D14 j2 D54 and D14 j4 D79), with D14 j4 D79 having the strongest effect. Interestingly, while D14 is strongly stabilising with some contextual dependence when used as part of a junction module, D14 j2 D14, it becomes more universal with a weaker stabilising effect. Whereas when used as part of D14 j1 D14, it has no stabilising effect at all. The latter is due to the presence of a junction domain which hinders the packing of helices into tight conformations.

To explore this unusual feature of the D14 module further, we considered constructs consisting of *k* consecutive repeats of D14 modules followed by 6 *− k* consecutive repeats of D14 j1 D14 or D14 j2 D14. For each of these constructs, we compared the distributions of pairwise distances between all carbon alphas and normalised these to the largest distance in the smallest construct (i.e., six repeats of D14). We found that as the number of D14s decrease and the number of D14 j1 D14s increases, the distributions flatten out and a second prominent mode emerges (**Figure 6c**). This coincides with a less tight packing that ultimately leads to a less stable behaviour. In contrast, as the number of D14s decrease and the number of D14 j2 D14s increases, the mode increases but the variance of the distribution decreases, leading to more consistent packing and stable behaviour.

### Modelling multi-chain constructs

So far, we have only considered single-chain repeats. However, building more complex structures quickly becomes infeasible, as many structures are not reducible to a single-chain design. Moreover, due to the repetitive nature of the sequences, long repetitive DNA molecules can be difficult to synthesise and large repeat proteins can be expresses in low yield. One approach that is observed in nature, and commonly used by protein engineers to overcome this limitation is to employ multiple chains that physically interact to form a larger structure.

Tree-like structures bring together two or more single-chain repeat domains at branching points, which act like ‘hubs’ within the structure. To facilitate the design of tree-like structures using Dynamo, we extended it’s capabilities to allow for hub modules within the parts database. Each hub is able to connect together multiple chains of modules together, allowing them to act as branches in the overall tree structure.

To accommodate tree-like structures in our model, we exploited the fact that when two modules are coupled, they physically interact and are stuck together. In other words, they do not necessarily have to be part of a single chain. This allowed us to loosen our definition of a module to merely regions of a protein that can be reused in different designs. In reality, hubs could either be an expressible protein, or could emerge from the cross interactions between two or more linear chains of modules. In this second case, we offer the ability to subjectively define a region around the interaction as a module. In terms of the abstraction in the model, we make no distinction between either case and treat both similarly (i.e., we represent hubs in the same way as modules, but with that added ability of being able to connect to more than two modules). With this extension, we can represent an arbitrary tree-like structure using a set of modules and a list of connections for each.

In order to predict the dynamics of tree-like structures, we exploited the fact that once represented using our model, the tree-like structure naturally defines a Bayesian network. We can therefore arbitrarily select an anchor module relative to which movement of other modules will be predicted, and then propagate movements using Eq. (1) for all branches emanating from the selected anchor module. The additional assumption underlying this approach is that the branches downstream of a hub do not physically interact with each other, which is valid when the ends of the branches are sufficiently separated.

To test this functionality, we designed a star-like multi-chain protein where a D4 C4 G1 hub module is connected to four independent chains of four D4 modules (**Figure 7a**,**b**). We then predicted the structural dynamics for two different anchoring points at the central hub and at the end of one of the arms. As expected, this showed that anchoring at the central hub reduced the overall movements that could be achieved by the arms, while anchoring a single arm allowed for larger arm movements (**Figure 7c**).

**Figure 7:**
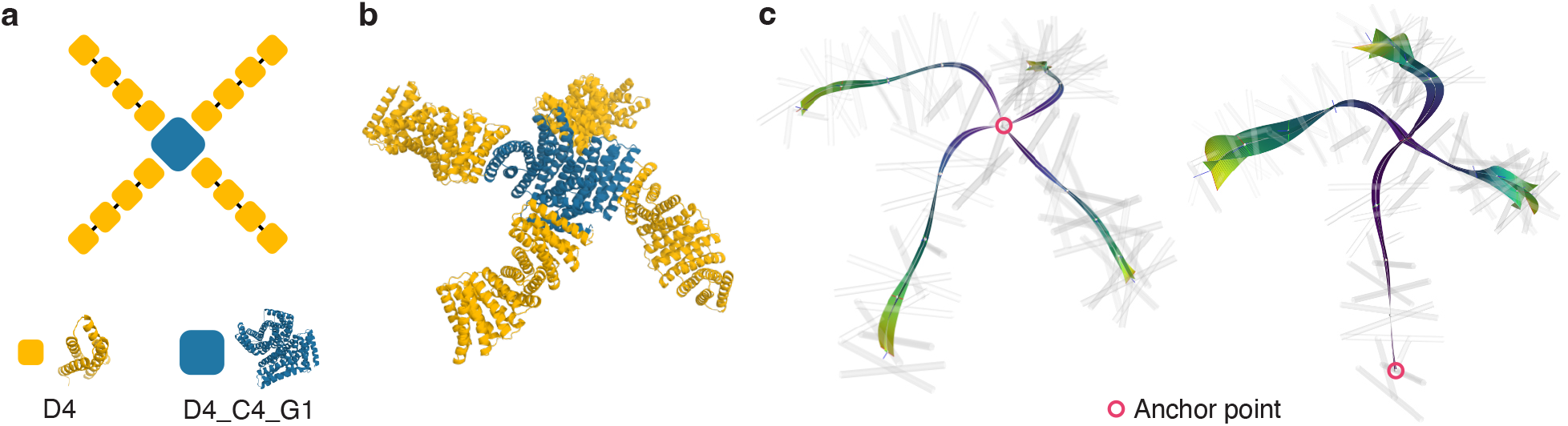
Estimating dynamics of a multi-chain tetrameter. (**a**) A schematic representation of a multi-chain construct and the two distinct modules used in the construct. (**b**) Molecular visualisation of the multi-chain construct. We use D4 C4 G1 hub with four D4 modules connected to each of the four arms of the hub. (**c**) A visualisation of the structural dynamics predicted using our model with two different anchor points (red circle).

## DISCUSSION

In this study, we developed a coarse-grained modelling approach to facilitate dynamics-driven repeat protein design. Our method successfully captured the essential features of modular protein dynamics and allowed for the exploration of their conformational space in a computationally efficient manner. For moderately sized proteins (4 or 6 modules long), the ability to calculate the conformational probability distributions and associated analyses in milliseconds on a standard desktop computer allowed us to exhaustively explore the structural dynamics for all possible designs, covering over 65,000 variants. The ability to provide such extensive coverage in protein design space enabled us to better understand how our design space covers particular features of interest, e.g., curvature and movement (**Figure 4**), and unravel the role that individual modules play in supporting the flexibility/rigidity of a resultant protein (**Figure 6**). Furthermore, the generality of our approach is not limited to single-chain repeat proteins. We show that a simple extension enables the prediction of structural dynamics of multi-chain, tree-like modular proteins (**Figure 7**) and the underlying mathematical model can accommodate any modular component for which samples of structural conformations can be gathered. The current work was focused on a library of compatible alpha helical modules, purely because of their availability, but, as more modular designs become available, our method can be apply to any modular system, even beyond proteins. Similarly, any dataset capturing population dynamics, either experimentally obtained or generated through simulations, could be used for the description and analysis of modular systems.

A major advantage of our approach is that it allows us to capture the inherent flexibility of repeat proteins. Modular repeat proteins often exhibit structural plasticity, allowing them to adapt and interact with different ligands or partners. By evaluating the dynamics of repeat proteins using our coarse-grained model, we are able to observe conformational changes and fluctuations in protein structure. These insights provide valuable information about the flexibility of different regions within the repeat protein and how they might contribute to its function. Understanding the flexibility of repeat proteins is also crucial for designing proteins with adjustable properties or for engineering proteins that can undergo conformational changes (e.g., upon binding to specific targets).

Our approach also revealed the multi-stability of many repeat proteins (**Figure 5**), which is a desirable property in many applications. Multi-stability refers to the ability of a protein to adopt multiple stable conformations or functional states. By exploring the conformational space of repeat proteins, we identified distinct energy minima corresponding to different conformations. This finding suggests that repeat proteins can exist in alternative stable states, potentially enabling them to switch between different functional states or adopt different binding configurations. Exploiting the multi-stability of repeat proteins opens up new opportunities for designing protein-based switches, sensors, or molecular machines with programmable functionalities. Our feature extraction method for identifying multi-stable features could be further refined to allow for greater specificity.

With the advent of AlphaFold ^44^, machine learning (ML) and generative artificial intelligence (AI) have become commonplace in protein design workflows ^45,46^. While these approaches offer unprecedented accuracy in the prediction of protein structure from sequence alone, their use for the prediction of protein dynamics has been limited ^47^. This stems in part from difficulties in generating the large training sets required, although there have been recent efforts to overcome these issues ^48,49^. A further challenge that remains is the high-computational cost of running ML models after training. While acceptable for small design spaces containing hundreds of possible designs, larger design spaces remain inaccessible due to the computational demands. An interesting future direction would be use ML to generate the conformational snapshots needed to parameterise the modules of the Dynamo model ^49^. This would then offer the means to blend ML predictions at the protein module level, with Dynamo’s efficiency in combining that data at the level of large single-chain repeat proteins or multi-protein assemblies.

The past decade has seen the design of *de novo* protein structures explode. Looking forward, the next frontier will be the design of protein dynamics and the push towards implementing complex molecular functions that require carefully choreographed structural changes over time. Tools like Dynamo will be crucial for accelerating our ability to practically explore the dynamics of repeat proteins and modular biological structures, supporting steps towards this goal.

## METHODS

### Repeat protein library

We use a reduced subset of an existing repeat protein library ^38,39^ consisting of 11 homo-modules, and 23 junction modules that consists of two homo-modules interfaced together with a junction modules. The repeat protein library can be used as part of the Elfin ^37^ tool or directly from the data repository (i.e., https://github.com/Parmeggiani-Lab/elfin-data) where atomistic position data for all of the modules are stored as PDB files in the compressed tarball `pdb aligned.tar.bz2`.

### Coarse-grained representation

Given that an atomic description is far too detailed for our purposes, it was important to find a simpler representation that is computationally tractable. An option would be to use the coordinates of the *C*_*α*_ atoms, and it would be possible to move to this level of detail at some point in the future, but for this work we find an even coarser description of secondary structures works well. Specifically, we use the STRIDE algorithm ^50,51^ as it exploits dihedral angle information in addition to the hydrogen bonds. While STRIDE can find all the structures in each of the conformations, the start and end locations can vary in the order of a few residues. Thus, to compare each of the conformations, we take the intersection of the start and end locations among all of the conformations. In other words, for a given alpha-helix identified, the residues that are part of it are given by all of the residues that are common across all of the conformations that have been attributed to the same alpha-helix.

### Generating the dynamics database

To build the dynamics database we employed the Elfin software suite ^37^ to construct first all possible three module constructs. An exhaustive approach was used resulting in 644 repeat proteins. Each of these repeat proteins are single chains and they consist of three modules with capping repeats at both ends. Capping repeats are used to ensure solubility of the proteins when expressed, but in this work, they also prevented edge-dependent artifacts, such as opening of the terminal helices during relax. These proteins were further relaxed using the Cartesian relax protocol in the Rosetta relax application ^40,52,53^ to obtain a low energy reference structure with packed side chains. Using these reference structures, an additional round of relaxation was used to obtain 100 conformations given as atomistic positions in PDB files. For this latter relaxation the FastRelax protocol was used and no conformations were rejected once they were obtained. All Rosetta related tasks were performed using Rosetta version 3.9 and ‘ref2015’ score function for the relaxations.

### Abstract representation and rules of combination

Before we consider the dynamics of modules in a chain, we first construct a set of rules and axioms that are necessary to construct larger proteins. We start with the 2D representation before generalising to obtain the 3D representation.

We have to design a unique local representation and connection rules that result in a unique “deterministic” solution. Let *C* = {*𝒰*_1_, *· · ·, 𝒰*_1_} be the set of modules and *⊕* be a non-commutative operation to combine two modules. Suppose a module is defined in the following way, (a) a bounding box defined by a set of points that are measured relative to the centroid of mass (centroid). (b) a vector defining the location of the adjacent centroid w.r.t the current ***ν***_*n*_. (c) a vector connecting the current centre of mass to the previous, ***ν***_*p*_. With these properties, we define the operation *ℳ*_1_ ⊕ *ℳ*_2_ as the following. (1) Translate *ℳ*_2_ such that its centroid, ***c***_2_ = ***c***_1_ + ***ν***_*n*:1*→*2_. (2) Rotate, *ℳ*_2_ about its centroid such that ***ν***_*p*:1*→*2_ is parallel to ***ν***_*p*:1*→*2_. Notice that with just the operation (1), it is not sufficient to create a unique *ℳ*_3_ = *ℳ*_1_ ⊕ *ℳ*_2_ as *ℳ*_2_ can freely rotate about its centroid. Having the vector ***ν***_*p*_ with which to align ***ν***_*n*_ ensures that ***ν***_*n*_ can only point in a single direction, i.e. removes radially non-uniqueness.

Generalising to 3D, we find the following problem. While, *ℳ*_2_ and *ℳ*_1_ are aligned with respect to the centroid and the ***ν***_*n*_, we find that properties (a)-(c) and (1) and (2) are no longer sufficient to remove the problem of non-uniqueness. This is due to the fact that the module *ℳ*_2_ can freely rotate about the vector ***ν***_*n*:1*→*2_. The simplest way to amend this is by having another vector which is not parallel to the ***ν***_*n*_

### Defining units for database

For our particular protein database, we can distil all of these ideas from the previous section into a concise system. Note that for clarity, we use the term unit to define the mathematical representation of the protein, and reserve the term module to refer to the actual protein module.

From our database, each one of the 644 repeat proteins, defines a unique triplet of modules, (_*L*_*ℳ*_*R*_). Given that the position and movement of elements within a module depend on its context, i.e. neighbouring the modules *L* and *R*, it is necessary to define a module for each unique triplet of modules. In other words, our database of units has a size of 644, whereas the number of unique modules is only 34.

To define a unit, from a triplet _*L*_*C*_*R*_ repeat protein, we employ the following steps. We first separate the helices that belong to *L, C* and *R*. We then compute centroids, ***c***_*L*_, ***c***_*C*_, and ***c***_*R*_ for the modules, *L, C* and *R*, respectively. We define a reference, ***R***^(*L→C*)^ = [***e***_1_, ***e***_2_, ***e***_3_]. The vectors ***e***_*i*_ are normalised orthogonal vectors, with ***e***_1_ parallel to ***c***_*C*_ *−* ***c***_*L*_, and ***e***_2_ parallel to ***e***_1_ *×* ***h***_*L*_, where ***h***_*C*_ is the vector from the mean bottom of helices in module *C* to the mean top. Having define, ***R***^(*L→C*)^, perform a rigid body transformation by multiplying all points in *C* and *ℛ* with the inverse of ***R***^(*L→C*)^. We can then define another reference frame for ***R***^(*C→R*)^ following the same constraints as before but modules *C* and *R*. To connect with the module *R*, we define the vector ***ν***_*n*_ = ***c***_*R*_ *−* ***c***_*C*_. Lastly, we can define any other reference points within the module *C*, e.g. those of a bounding convex hull. In summary, we compute ***R***^(*C→R*)^, ***ν***_*n*_ and any bounding box points relative to the centroid of *C* and with the additional constraint that ***R***^(*L→C*)^ = ***I***.

With this abstract representation, we define the non-commutative operation ⊕, for *ℳ*_1_ ⊕ *ℳ*_2_ as the following. (1) We translate *ℳ*_2_ such that its centroid is at ***c***_1_ + ***ν***_*n*:1*→*2_. (2) Perform a rigid body transformation for all of the vectors and reference frame in *ℳ*_2_ by the ***R***^(*C→R*)^ reference frame of *ℳ*_1_. With these two rules, we can construct any valid chain of modules.

### Generalising the module abstraction to include dynamics

Having designed the module framework, we now move on to generalising the frame to capture the dynamics involved. Such a generalisation is relatively straightforward, one can define vectors within the framework, which includes the basis vectors in reference frames to be probabilistic. In other words, instead of using a single vector, we use a cloud of vectors given by a probability density function which we call probabilistic vectors.

Depending on which aspects are turned into a probabilistic vector, we can approximate the dynamics with increasing fidelity: 1. A static representation as defined in the previous sections; 2. The centroid vectors are probabilistic which captures bulk location. 3. The centroid vectors are probabilistic, and the reference frames connecting modules are probabilistic which together capture bulk location and bulk orientation. 4. The centroid vectors are probabilistic, and vectors to reference points are probabilistic. Captures bulk location orientation and location of reference points. For our analysis, we consider only the fourth case. It is important to note that the addition of two probabilistic vectors (p-vectors), is a convolution, while a rotation results in the rotation of the mean and covariance of the distributions.

To obtain these probability vectors from the conformations, we let *M*_*i*_ and ***c***_*i*_ be the number of points of interest and the location of the centroid for the *i*^th^ module. For each module, we then define a set of probability distributions given by

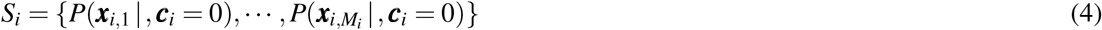

where ***x***_*i, j*_ is the *j*^th^ point of interest of the *i*^th^ module. In addition, to *S*_*i*_, we define the coupling distribution *P*(***c***_*i*+1_ | ***c***_*i*_ = 0) that describes the steady-state movement of the next module ***c***_*i*+1_ relative to the present one ***c***_*i*_, giving a total of *M*_*i*_ + 1 p-vectors.

In order to represent these p-vectors, an appropriate probability density estimation is required that satisfies a set of criteria: (i) it must have a parametric description for efficient storage; (ii) the density estimation must allow for rapid convolutions; (iii) the number of parameters must be “containable” so that it does not grow too large when we have many convolutions; and (iv) must be easily computable with arbitrary moments. These criteria are satisfied by a Gaussian mixture model. To fit a Gaussian mixture model with the appropriate number of components, we employed a *k*-fold validation protocol on the conformational data. We first train a Gaussian mixture with a different number of components, *n*, and use the likelihood estimate of the test set from the model to score the fit. We selected the number of components that gave the highest likelihood score.

### Flexibility score to assess protein rigidity

To assess the rigidity of a protein, we exploited the fact that this feature manifests itself in the model as centroids with distributions that have narrow variance. One can imagine an isosurface that expands outwards from a fixed centroid to the opposite which has the most movement. The larger the volume encapsulated by the surface the more flexible the protein and vice versa. To estimate this isosurface, for each centroid, we sampled from its Gaussian mixture density estimate giving three-dimensional points in space. We then projected these points onto a plane that is normal to the centroid backbone. Using this, we can fit a two-dimensional normal distribution to the points on the plane, from which an elliptical contour can be inferred that captures a given amount of the variance. In our case, we used the 95^th^ percentile, which is approximately two standard deviations from the mean. Connecting the ellipses with a linear interpolation gave us a pseudo-isosurface from which we then computed the volume enveloped. For convenience, we defined the cubic root of this volume as the flexibility score (*β*) so that the score scales linearly with the number of modules. Given two constructs, the more rigid example will have a narrower envelope of movement resulting in a lower isosurface volume and *β* score. Such flexibility scores provided a convenient way to compare the rigidity of different proteins.

### Computational tools

All computational simulations and analyses were run using Python version 3.11.4.

## DATA AVAILABILITY

The Dynamo package used to generate all the results for this article is split into two parts. The first part, called ‘dynamo’, is a native Python library built in Rust for evaluating the steady-state dynamics of large bio-molecular constructs. This library is not focused on modular repeat proteins and can be used for any modular structures. It is available at: https://github.com/seeralans/dynamo. The second part, called ‘dynamo-rp’, is a Python library for coarse-grained modelling of repeat proteins and is available at: https://github.com/seeralans/dynamo-rp.

### ACKNOWLEDGEMENTS

This work was primarily funded by UKRI grant BB/W012448/1 (to T.E.G. and F.P.) In addition, T.E.G. was supported by a Royal Society University Research Fellowship grants UF160357 and URF/R/221008, and a Turing Fellowship from The Alan Turing Institute under EPSRC grant EP/N510129/1. F.P. was supported by an EPSRC Early Career Fellowship grant EP/S017542/1. The funders had no role in study design, data collection and analysis, decision to publish or preparation of the manuscript.

## AUTHOR CONTRIBUTIONS

T.E.G. and F.P. conceived the study, supervised the work, helped establish the methodology, aided with the interpretation of the results, and edited the manuscript. S.S. developed the methodology, implemented the approach, carried out all experiments and analyses, and wrote the manuscript. T.E.N. provided the initial dataset.

